## Supplemental Information for "Sex- and context-dependent effects of acute isolation on vocal and non-vocal social behaviors in mice"

### S1 Fig. Effects of acute isolation on latency to first USV in same-sex and opposite-sex interactions

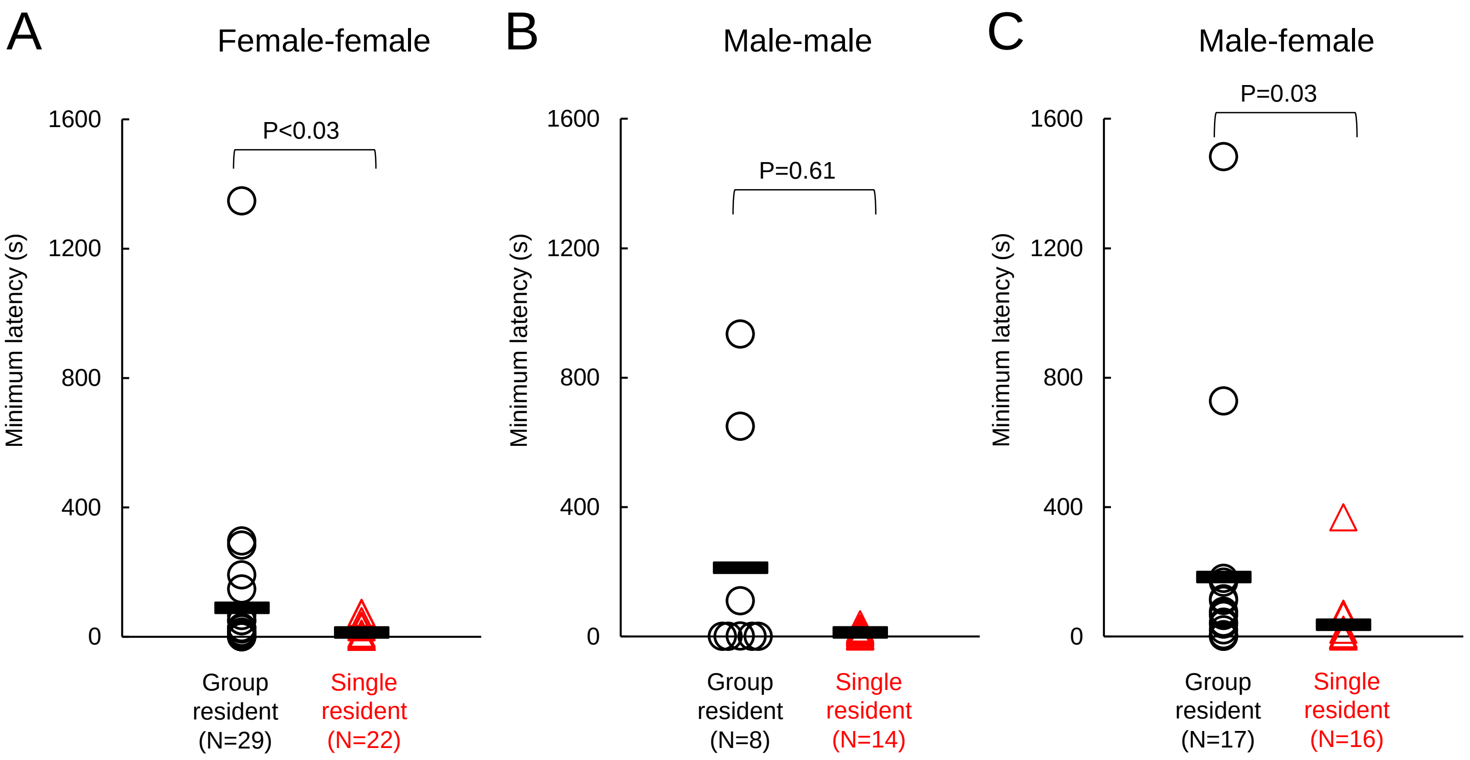


**S1 Fig. Effects of acute isolation on latency to first USV in same-sex and opposite-sex interactions.** Latency to the first recorded USV is shown for female-female social encounters (A), male-male encounters (B), and male-female encounters (C). Trials with 0 USVs are excluded.

S2 Fig (1/3). Female-female ethograms, group-housed residents

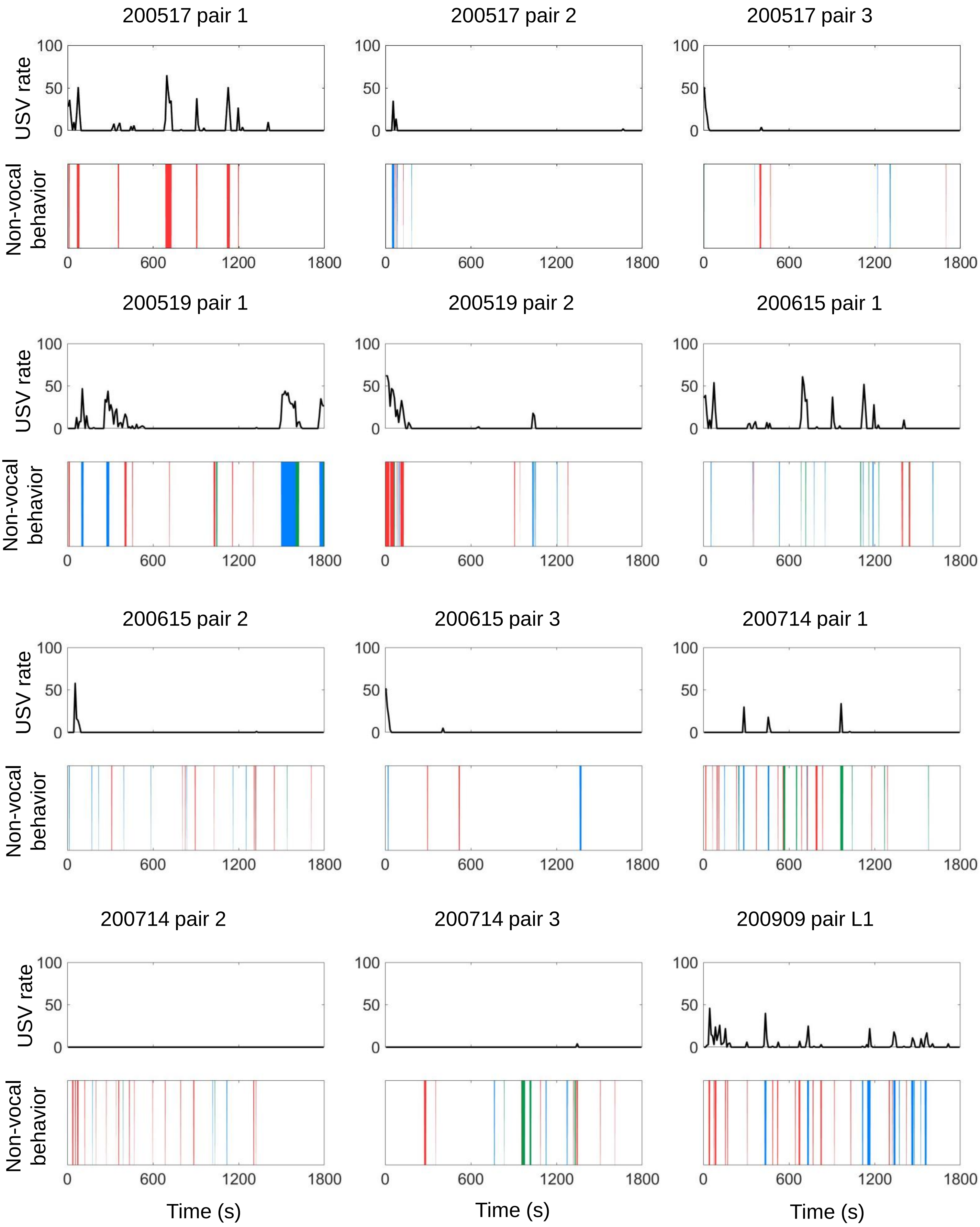

### S2 Fig (2/3). Female-female ethograms, group-housed residents

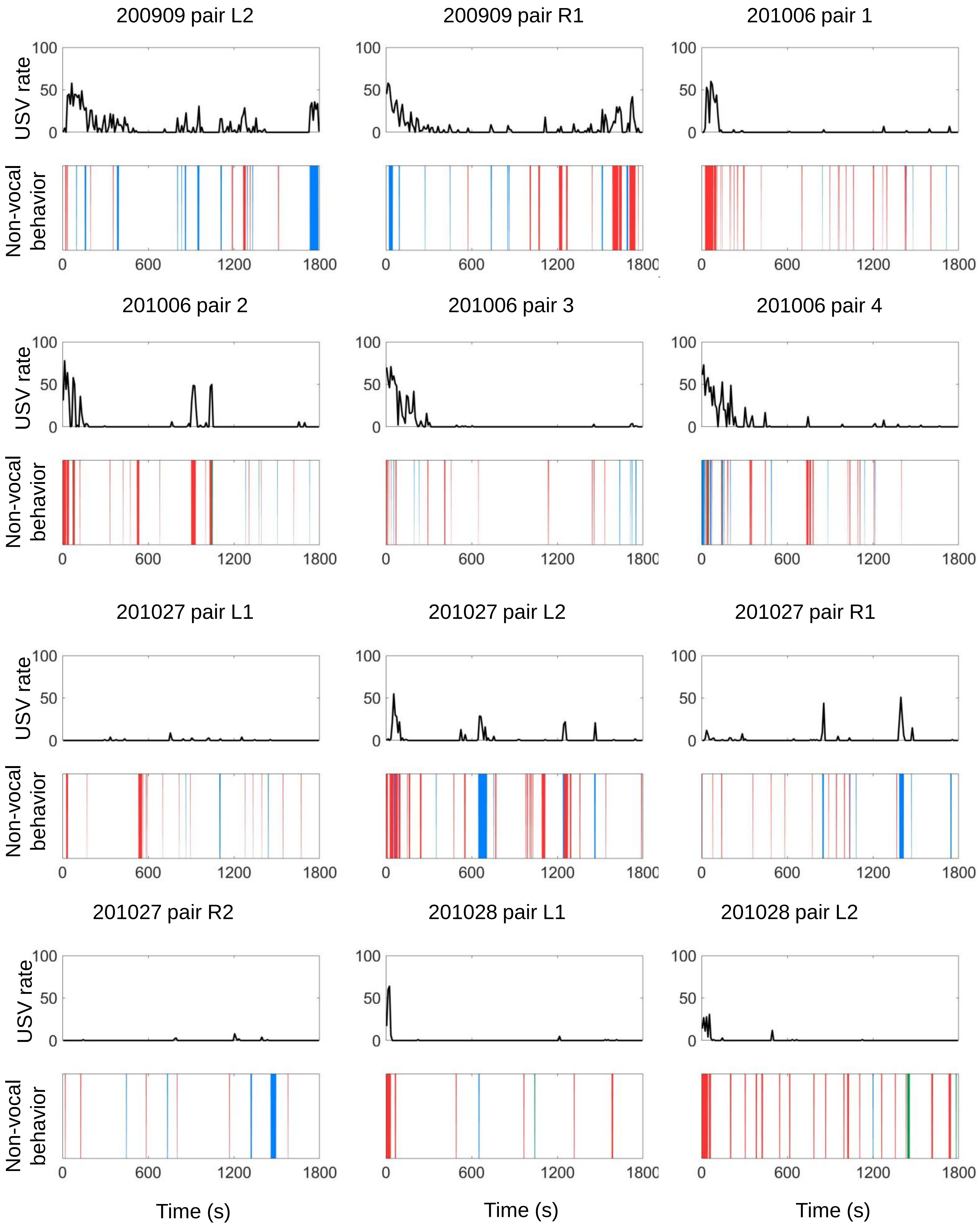

### S2 Fig (3/3). Female-female ethograms, group-housed residents

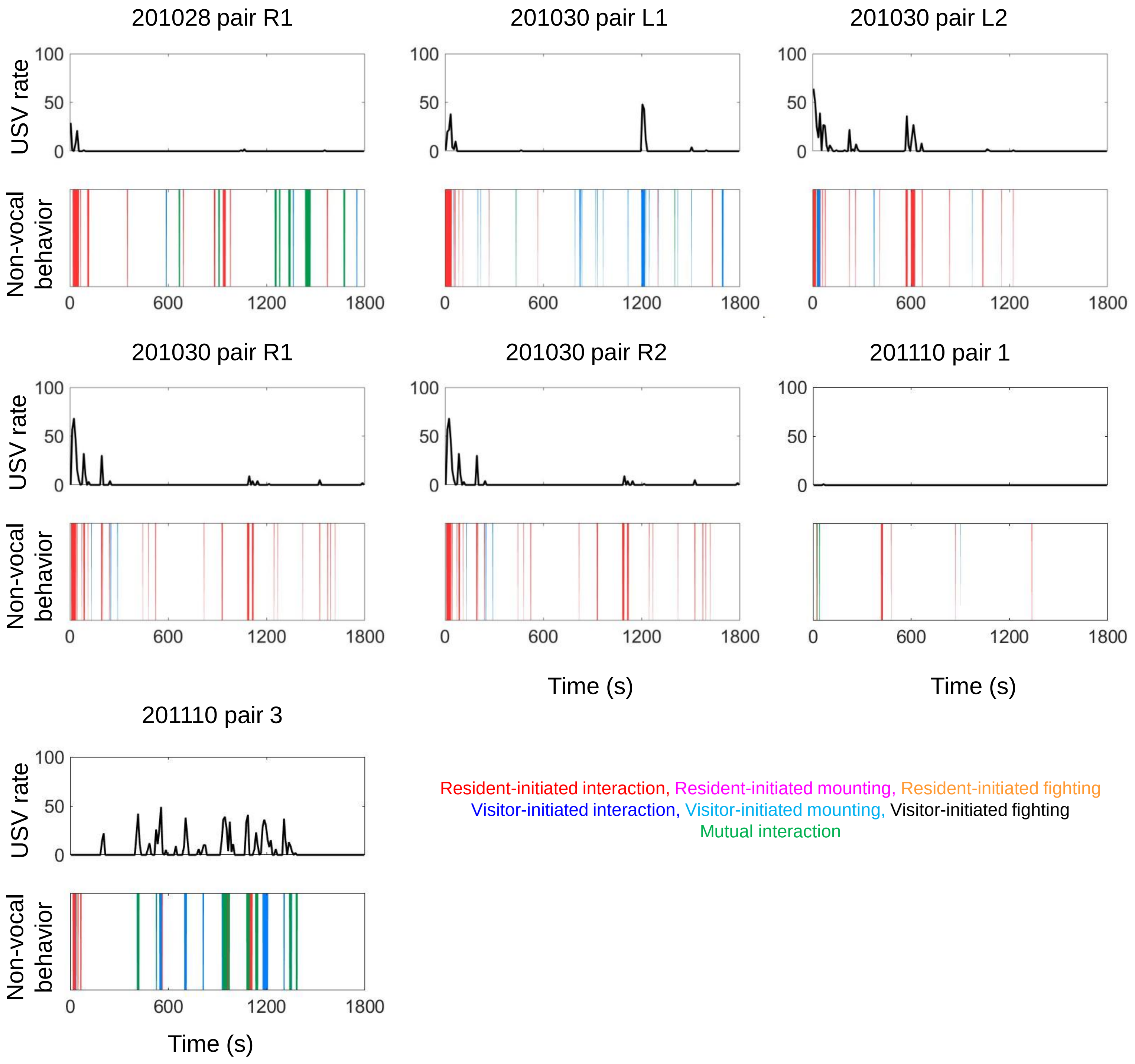

**S2 Fig. Female-female ethograms, group-housed residents.** Ethograms are shown for each female-female trial with a group-housed resident. The top half of each plot shows USV rate over time (total USVs in each 10s-long bin), and the bottom half of each plot shows the occurrence of different non-vocal social behaviors over time. Red, resident-initiated interaction; magenta, resident-initiated mounting; orange, resident-initiated fighting; blue, visitor-initiated interaction; cyan, visitor-initiated mounting; black, visitor-initiated fighting; green, mutual interaction; white, not interacting.

### S3 Fig (1/2). Female-female ethograms, single-housed resident

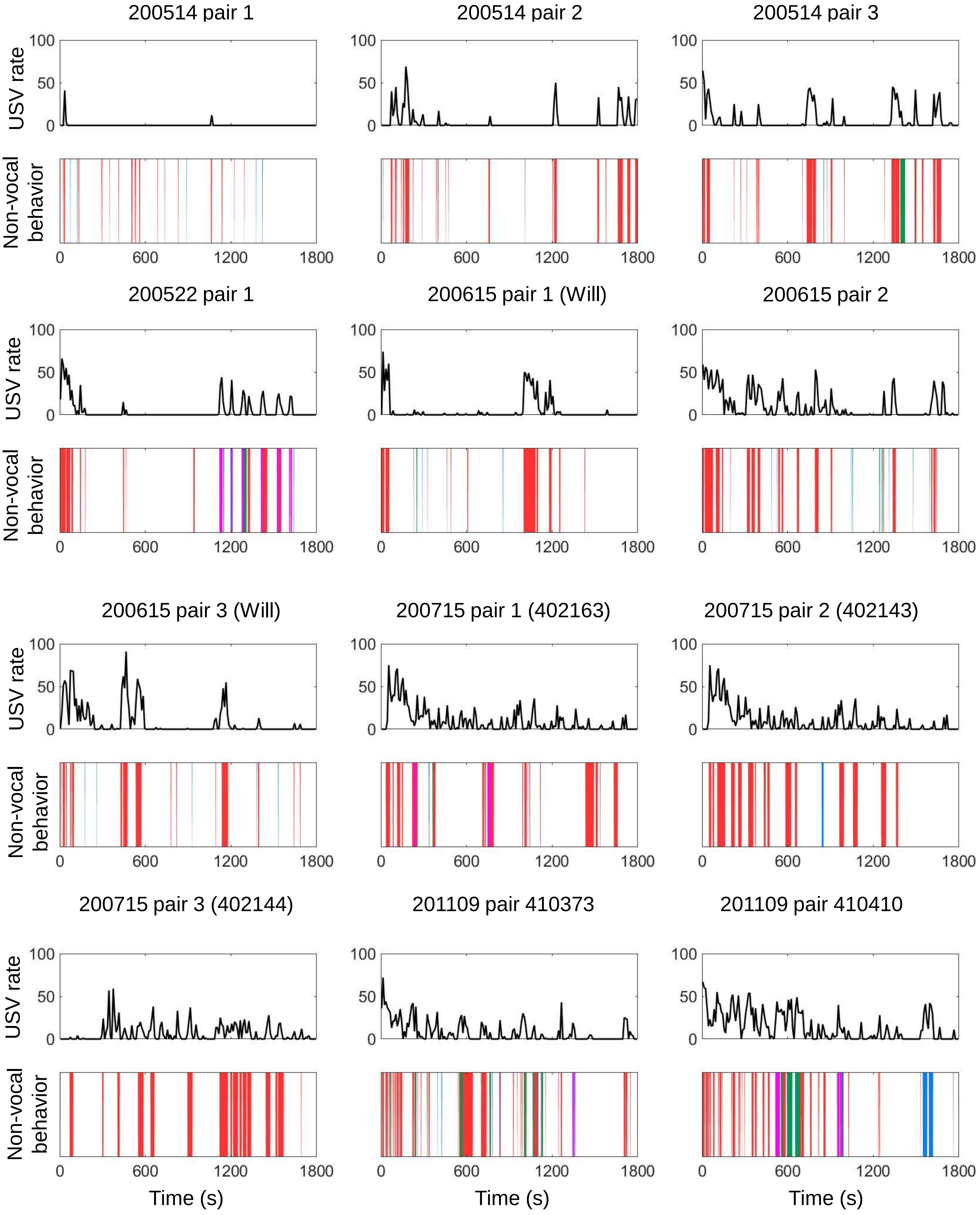

### S3 Fig (2/2). Female-female ethograms, single-housed resident

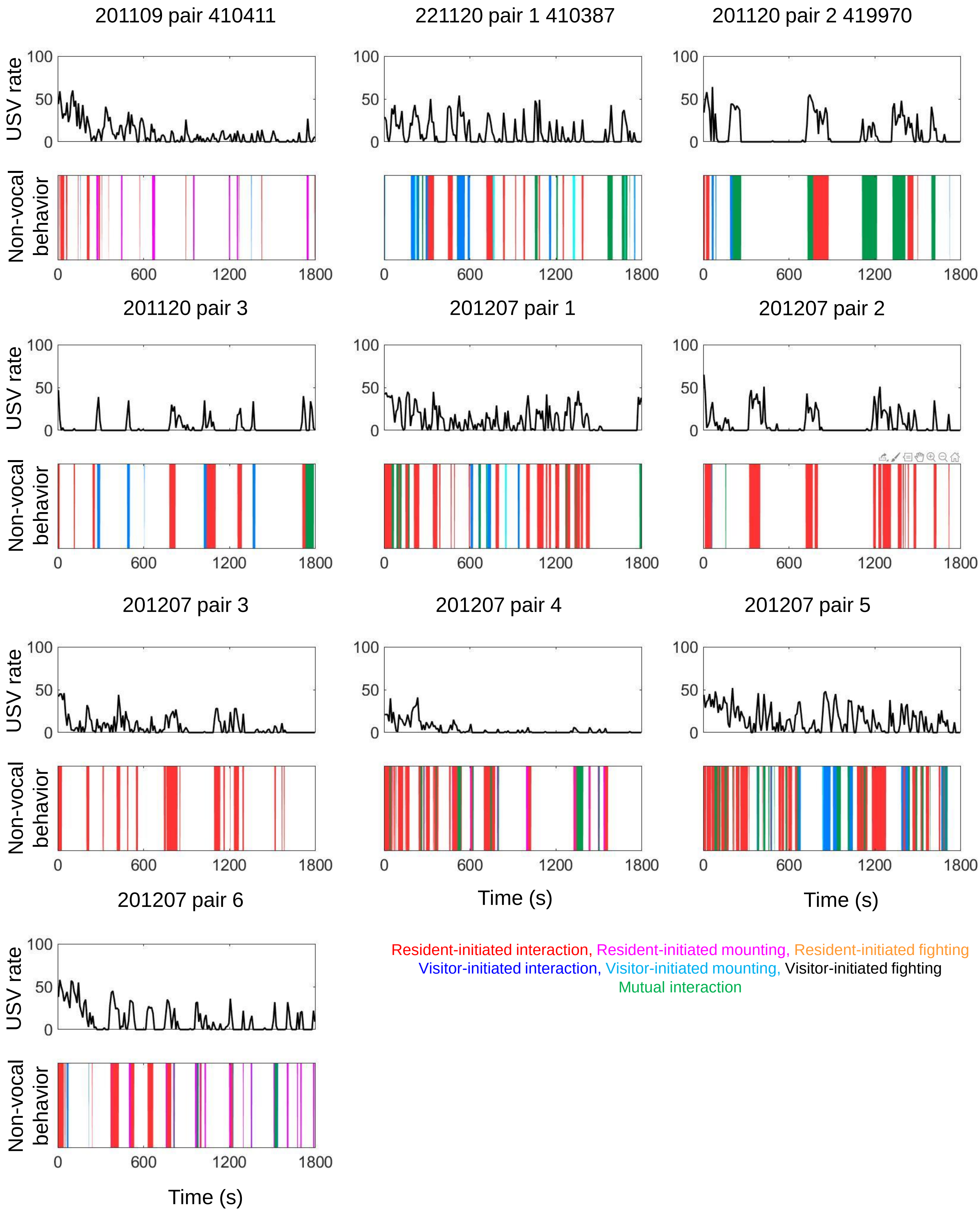

**S3 Fig. Female-female ethograms, single-housed residents.** Ethograms are shown for each female-female trial with a single-housed resident. The top half of each plot shows USV rate over time (total USVs in each 10s-long bin), and the bottom half of each plot shows the occurrence of different non-vocal social behaviors over time. Red, resident-initiated interaction; magenta, resident-initiated mounting; orange, resident-initiated fighting; blue, visitor-initiated interaction; cyan, visitor-initiated mounting; black, visitor-initiated fighting; green, mutual interaction; white, not interacting.

S4 Fig (1/2). Male-male ethograms, group-housed resident

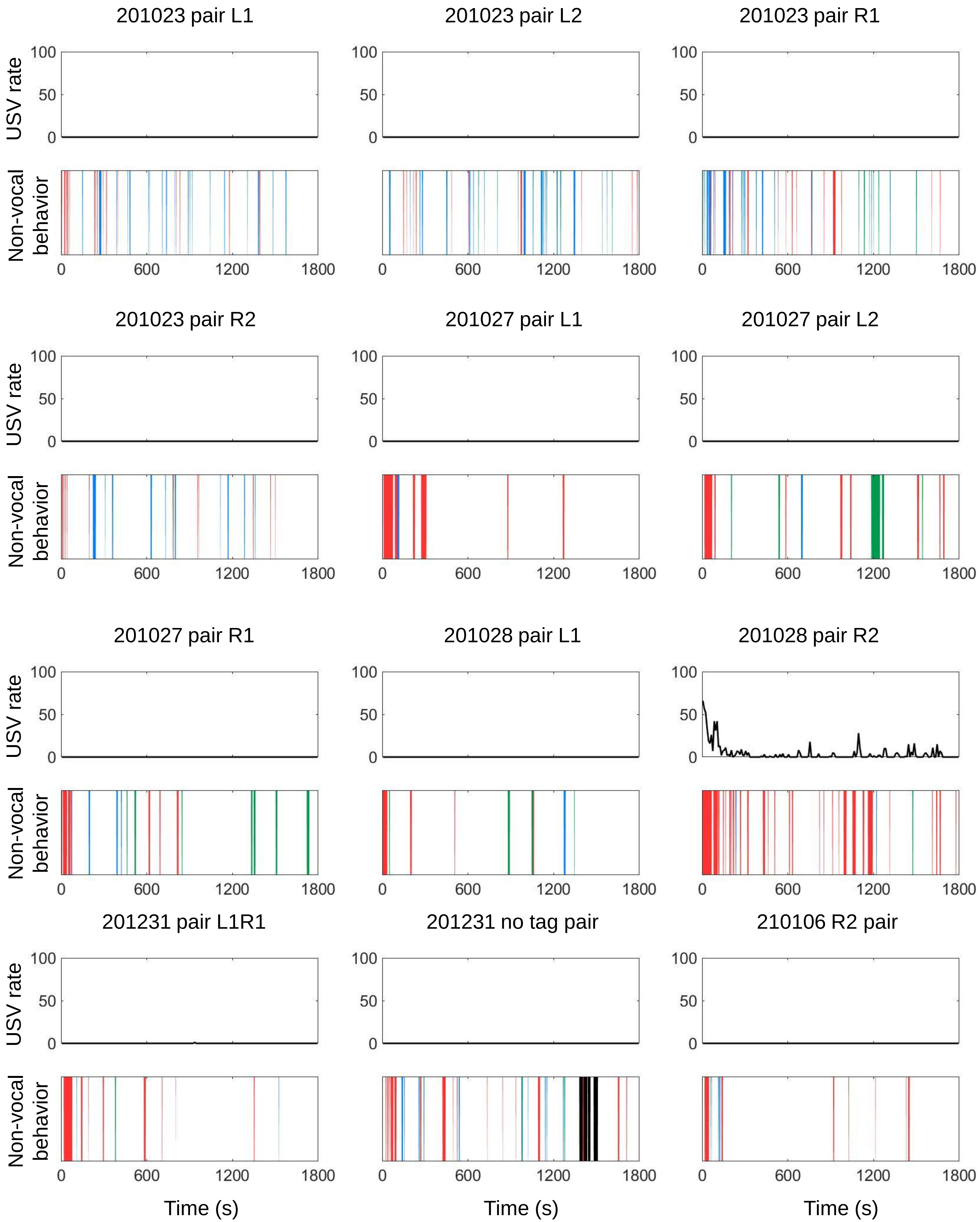

### S4 Fig (2/2). Male-male ethograms, group-housed resident

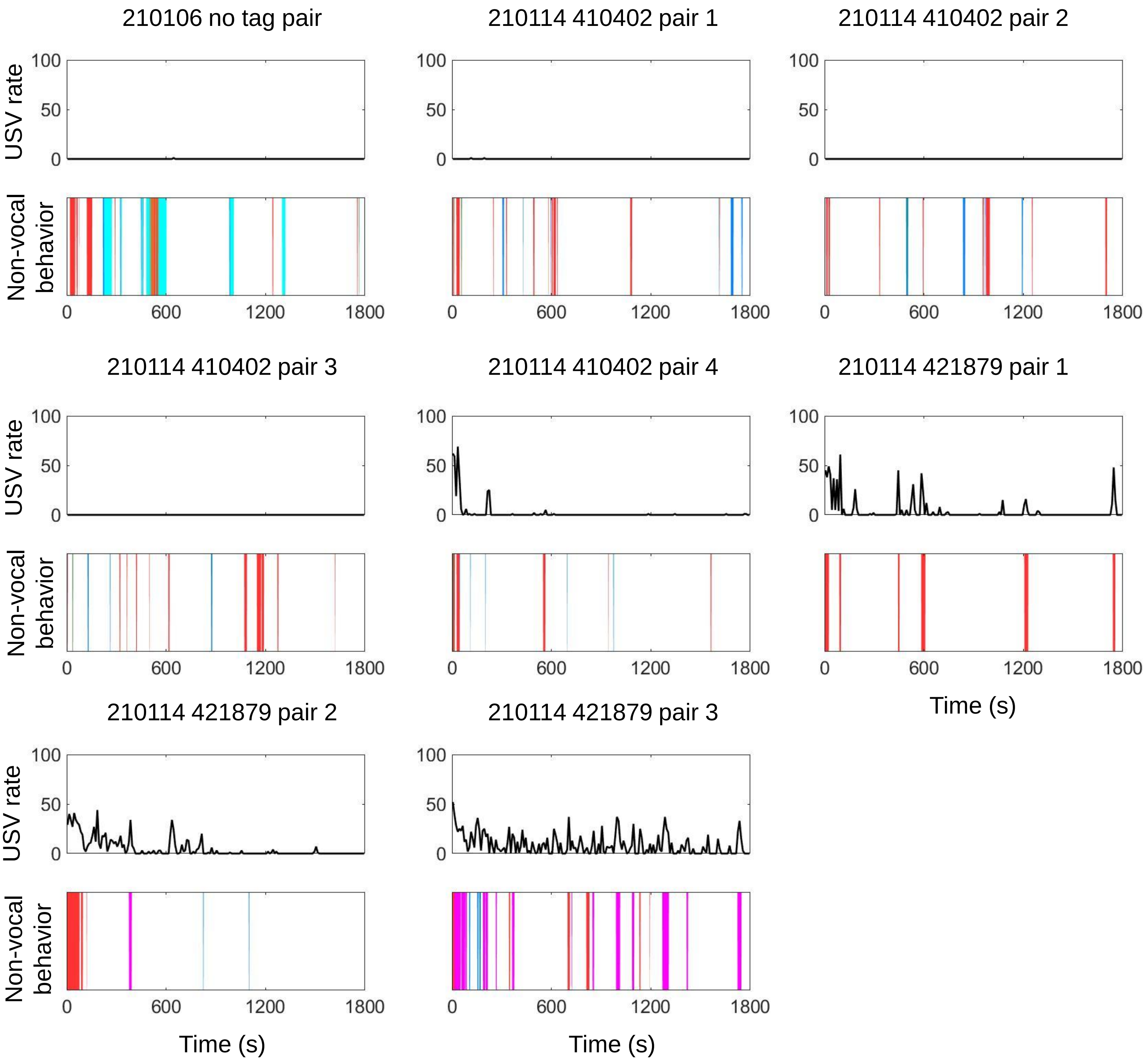


Resident-initiated interaction, Resident-initiated mounting, Resident-initiated fighting
Intruder-initiated interaction, Intruder-initiated mounting, Intruder-initiated fighting
Mutual interaction

**S4 Fig. Male-male ethograms, group-housed residents.** Ethograms are shown for each male-male trial with a group-housed resident. The top half of each plot shows USV rate over time (total USVs in each 10s-long bin), and the bottom half of each plot shows the occurrence of different non-vocal social behaviors over time. Red, resident-initiated interaction; magenta, resident-initiated mounting; orange, resident-initiated fighting; blue, visitor-initiated interaction; cyan, visitor-initiated mounting; black, visitor-initiated fighting; green, mutual interaction; white, not interacting.

#### S5 Fig. (1/2). Male-male ethograms, single-housed resident

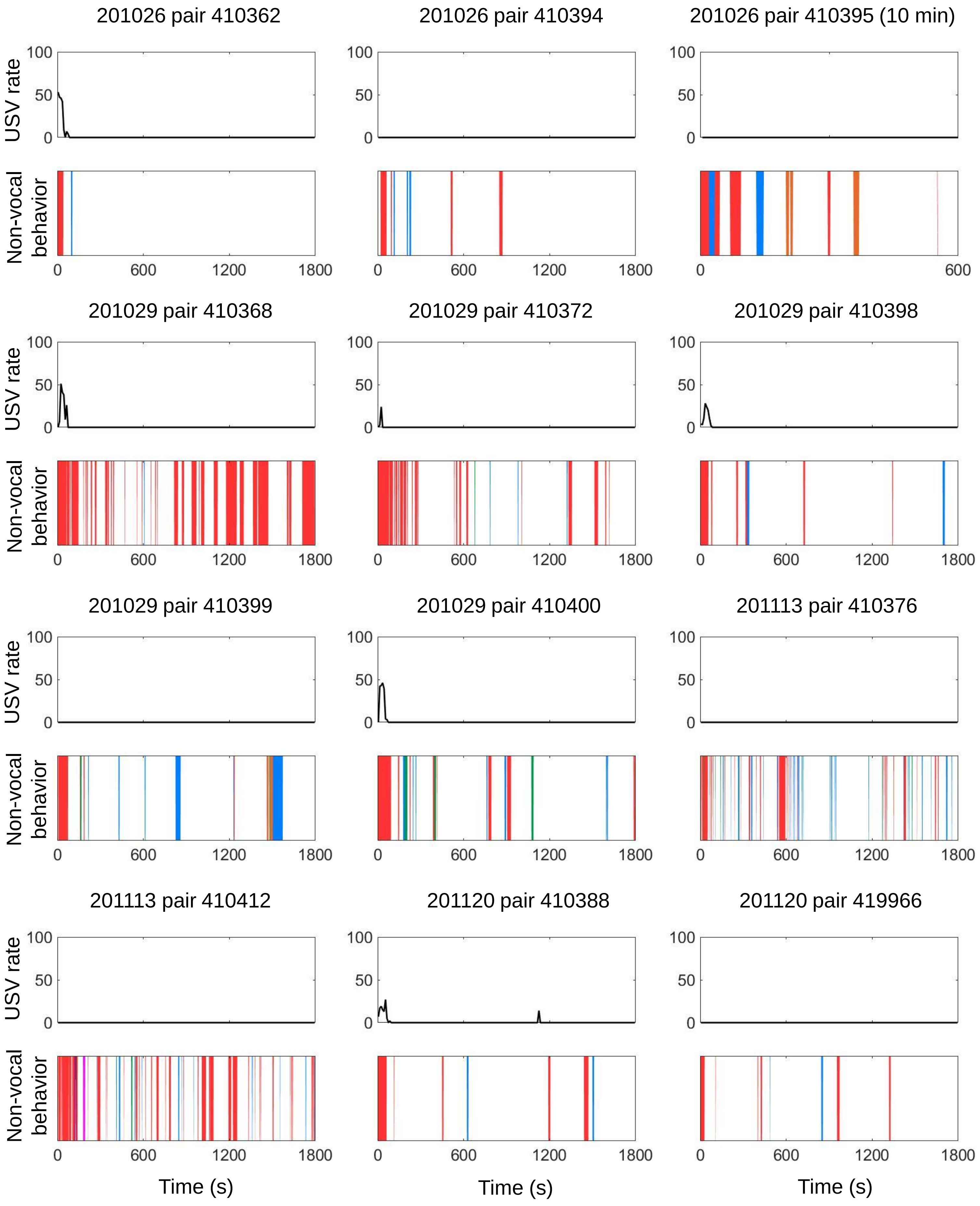

### S5 Fig (2/2). Male-male ethograms, single-housed resident

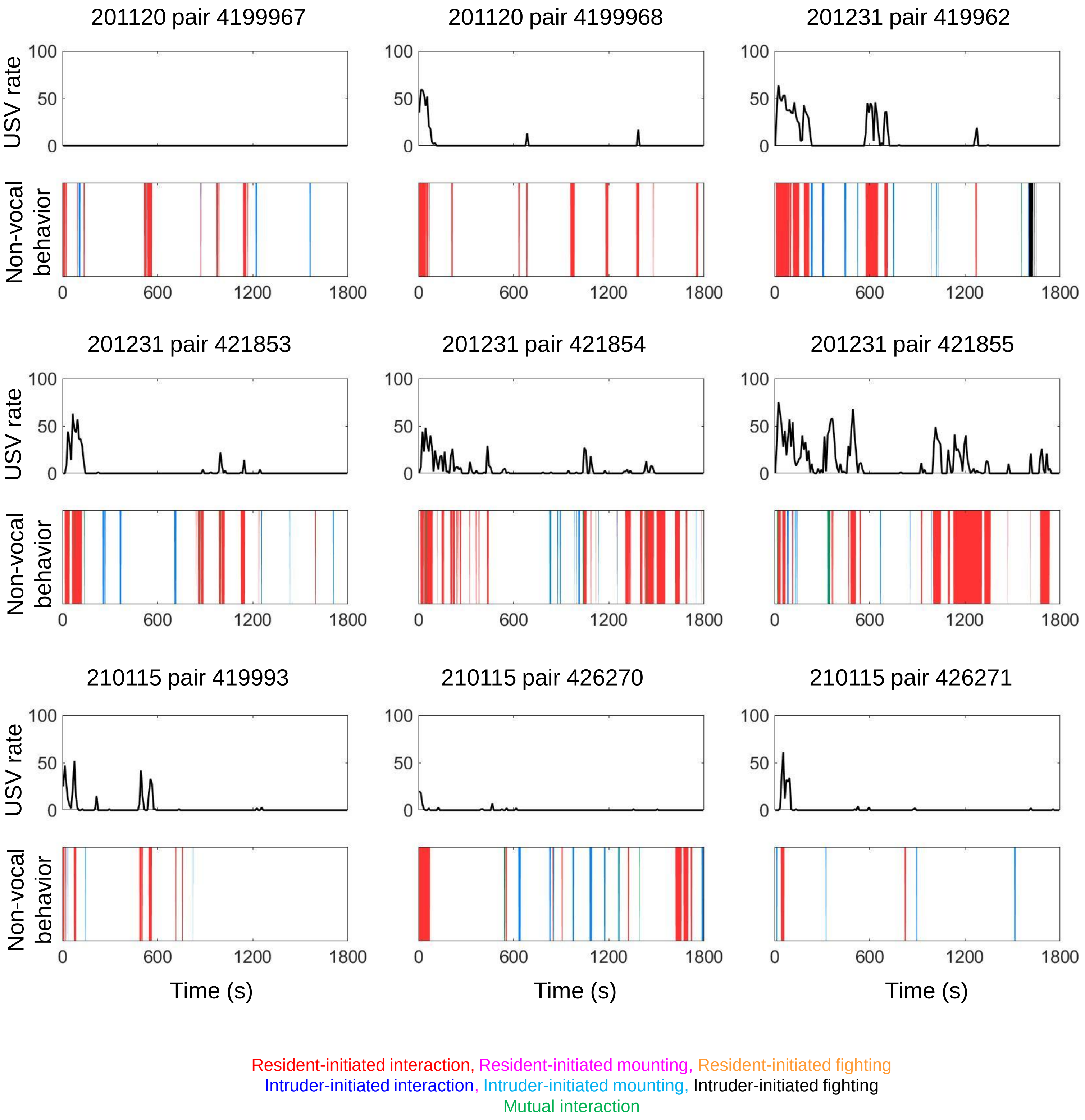

**S5 Fig. Male-male ethograms, single-housed residents.** Ethograms are shown for each male-male trial with a single-housed resident. The top half of each plot shows USV rate over time (total USVs in each 10s-long bin), and the bottom half of each plot shows the occurrence of different non-vocal social behaviors over time. Red, resident-initiated interaction; magenta, resident-initiated mounting; orange, resident-initiated fighting; blue, visitor-initiated interaction; cyan, visitor-initiated mounting; black, visitor-initiated fighting; green, mutual interaction; white, not interacting.

### S6 Fig (1/2). Male-female ethograms, group-housed resident

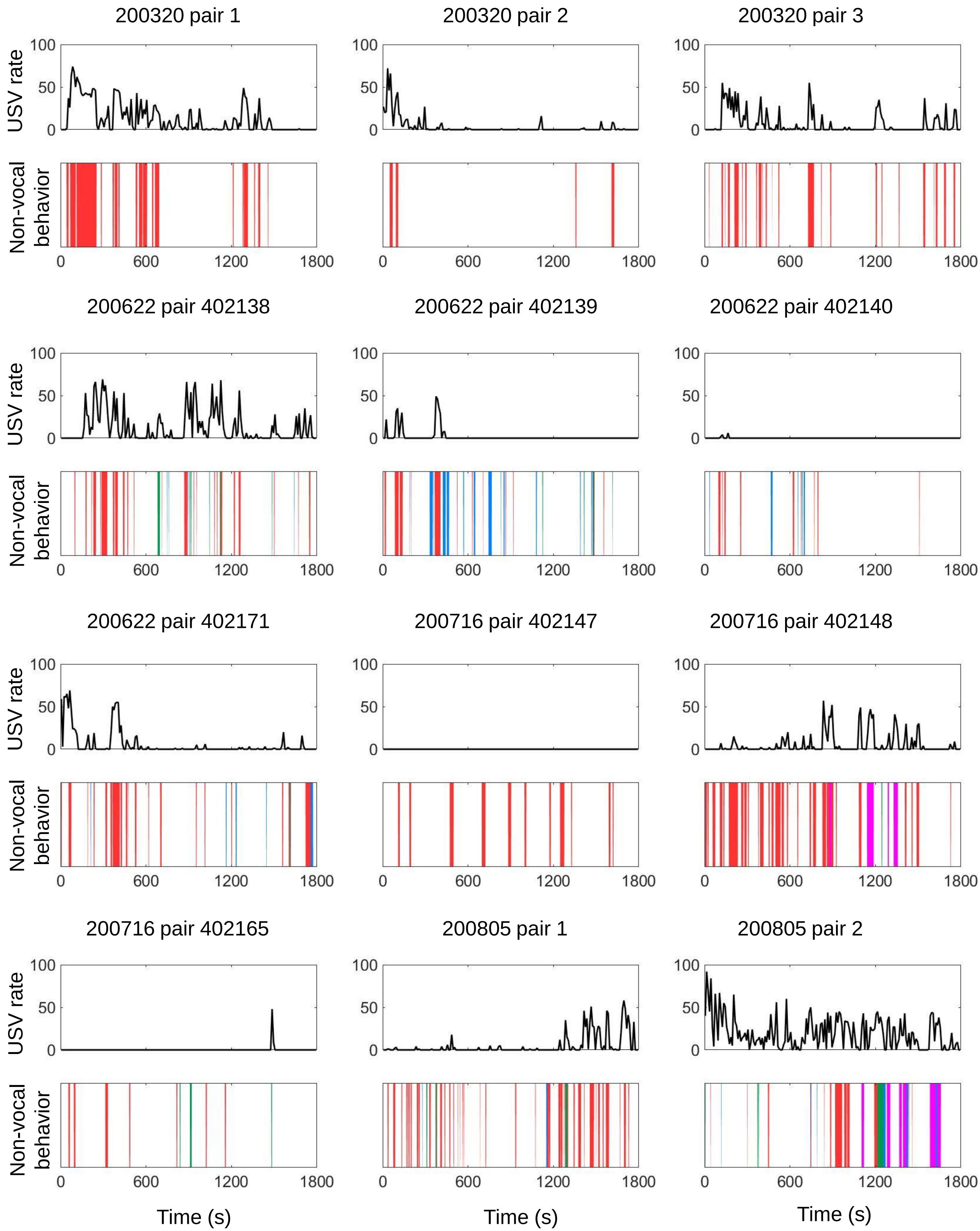

### S6 Fig (2/2). Male-female ethograms, group-housed resident

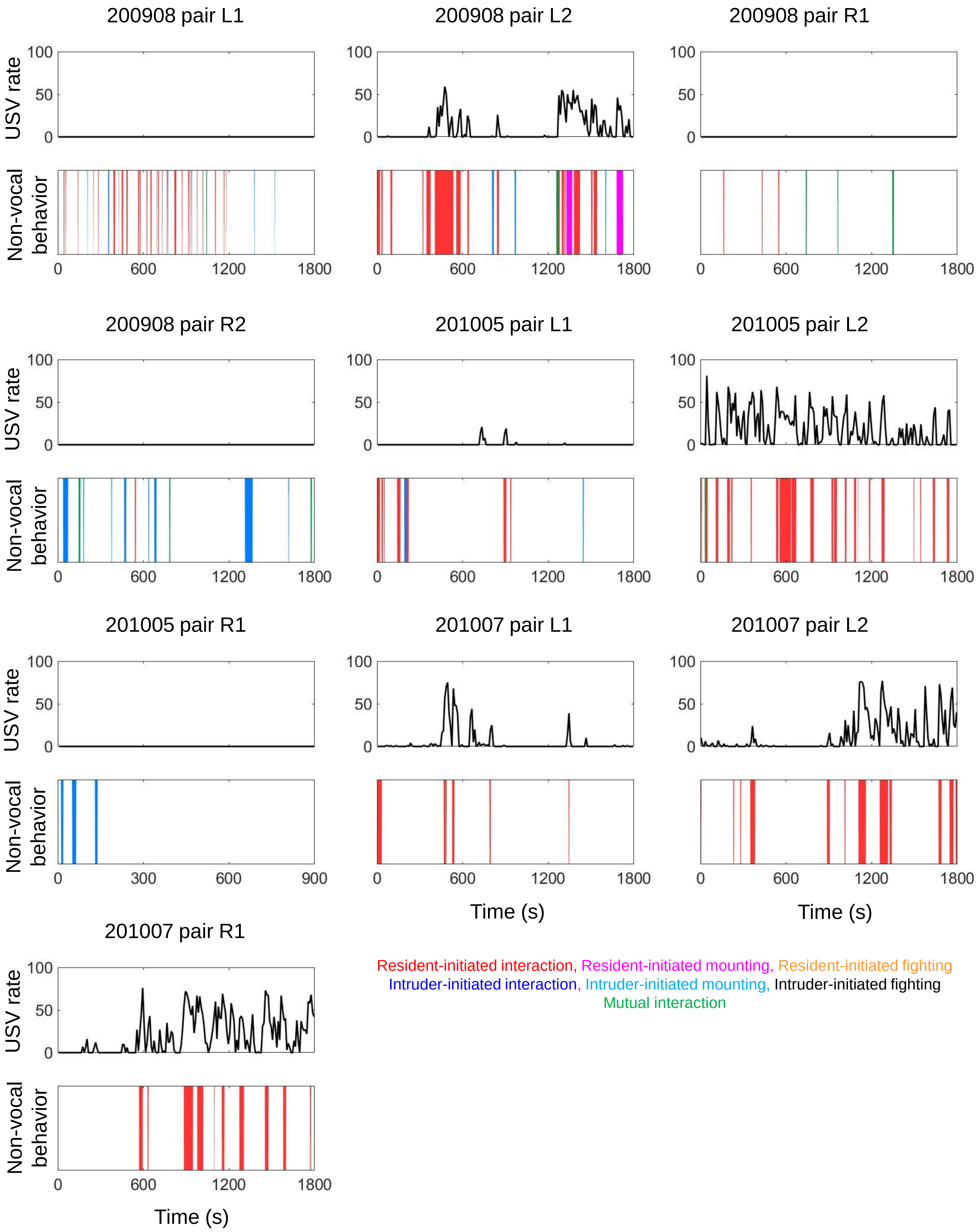

**S6 Fig. Male-female ethograms, group-housed residents.** Ethograms are shown for each male-female trial with a group-housed resident. The top half of each plot shows USV rate over time (total USVs in each 10s-long bin), and the bottom half of each plot shows the occurrence of different non-vocal social behaviors over time. Red, resident-initiated interaction; magenta, resident-initiated mounting; orange, resident-initiated fighting; blue, visitor-initiated interaction; cyan, visitor-initiated mounting; black, visitor-initiated fighting; green, mutual interaction; white, not interacting.

### S7 Fig (1/2). Male-female ethograms, single-housed resident

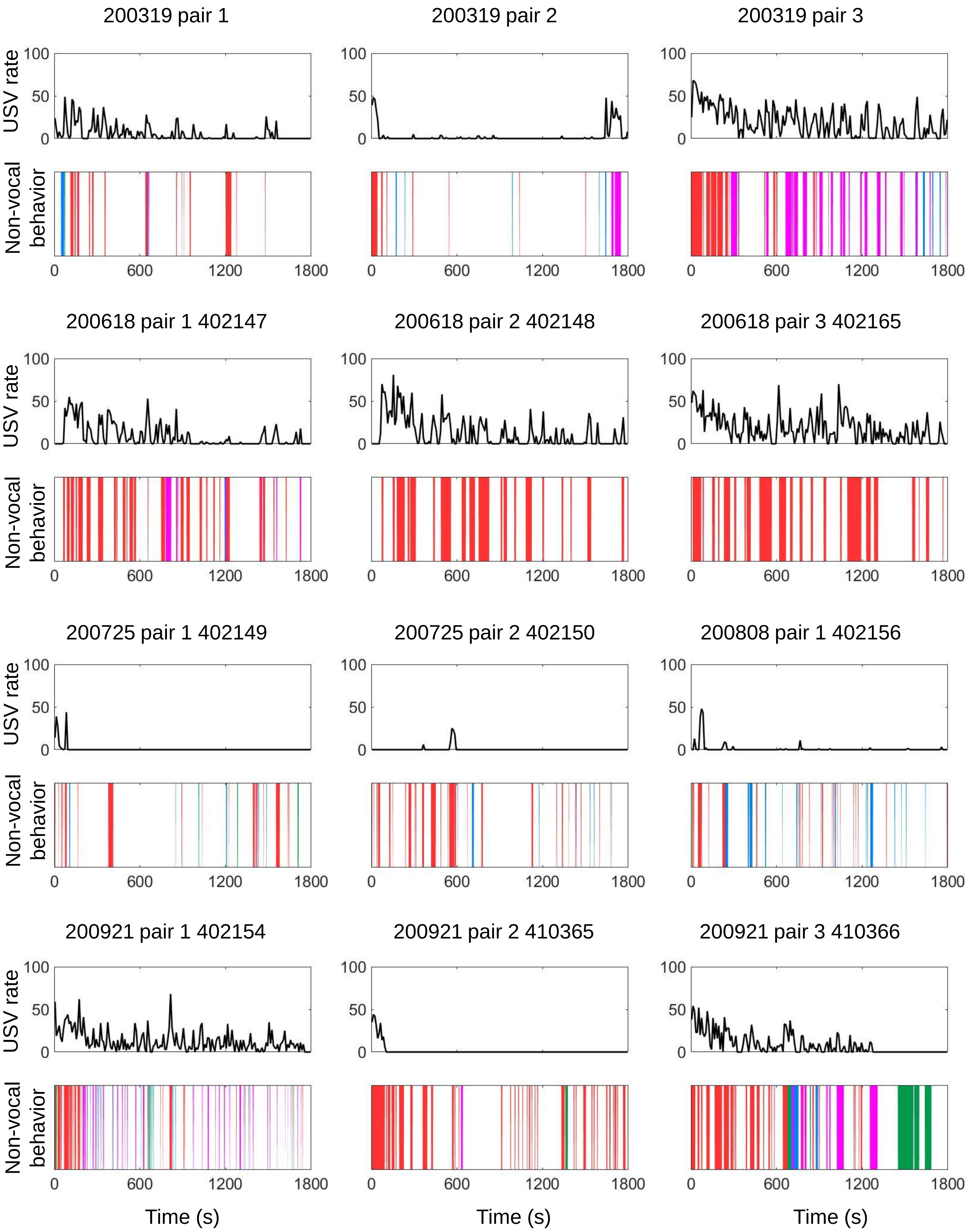

### S7 Fig (2/2). Male-female ethograms, single-housed resident

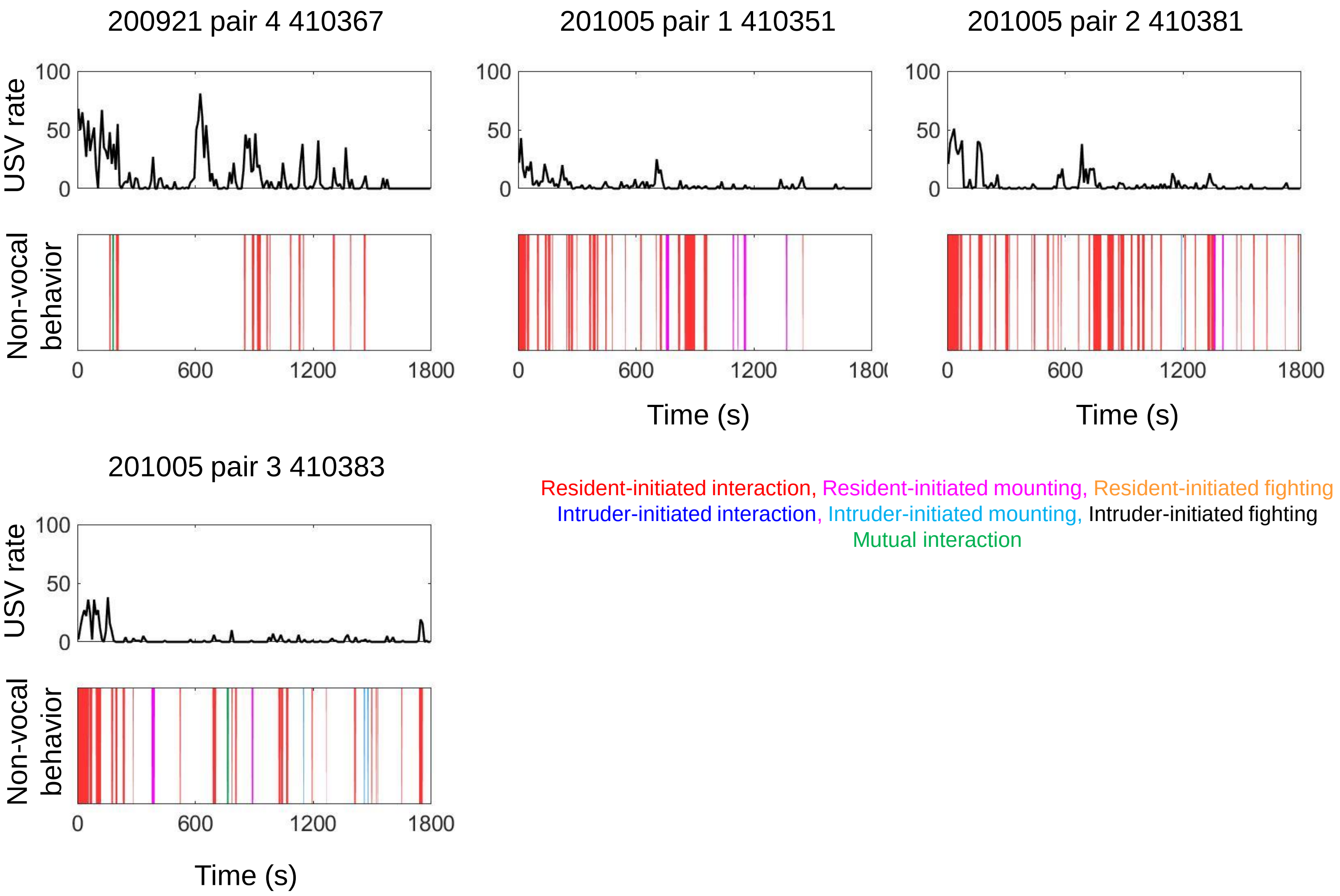

**S7 Fig. Male-female ethograms, single-housed residents.** Ethograms are shown for each male-female trial with a single-housed resident. The top half of each plot shows USV rate over time (total USVs in each 10s-long bin), and the bottom half of each plot shows the occurrence of different non-vocal social behaviors over time. Red, resident-initiated interaction; magenta, resident-initiated mounting; orange, resident-initiated fighting; blue, visitor-initiated interaction; cyan, visitor-initiated mounting; black, visitor-initiated fighting; green, mutual interaction; white, not interacting.

**S1 Table. Statistical Summary**

| Figure Number | Statistical Test | P value | Description |
|---|---|---|---|
| Fig 1A, left | Mann Whitney | <0.0001 | USV counts, female-female pairs |
| Fig 1B, left | Mann Whitney | 0.15 | USV counts, male-male pairs |
| Fig 1C, left | Mann Whitney | 0.44 | USV counts, male-female pairs |
| Fig S1A | Mann Whitney | 0.03 | USV latency, female-female pairs |
| Fig S1B | Mann Whitney | 0.61 | USV latency, male-male pairs |
| Fig S1C | Mann Whitney | 0.03 | USV latency, male-female pairs |
| text only | z-test for 2 independent proportions | 0.02 | proportion of male-male trials with >25 USVs |
| Fig 2A, left | Mann Whitney | <0.001 | time spent interacting, female-female pairs |
| Fig 2A, middle | Mann Whitney | 0.08 | time spent interacting, male-male pairs |
| Fig 2A, right | Mann Whitney | 0.08 | time spent interacting, male-female pairs |
| Fig. 2B | two-way ANOVA, repeated measures on one factor<br>(A, B, and interaction) | <0.0001<0.0001<0.0001 | differences in female-female non-vocal social behaviors |
| | | <0.0001<0.0001<0.0001 | differences in male-male non-vocal social behaviors |
| | | >0.05<br><0.001<br>>0.05 | differences in male-female non-vocal social behaviors |

| | | | |
|---|---|---|---|
| Fig. 2B | Post-hoc tests (resident-initiated, visitor-initiated, mutual) | <0.0001 0.18. 0.06 | differences in female-female non-vocal social behaviors |
| | | 0.003 0.25 0.26 | differences in male-male non-vocal social behaviors |
| text and Fig. 2C | z-test for 2 independent proportions | p<0.001 | proportion of female-female trials with mounting |
| | | p=0.27 | proportion of male-male trials with mounting |
| | | p=0.005 | proportion of male-female trials with mounting |
| text only | z-test for 2 independent proportions | 0.41 | proportion of male-male trials with fights |
| Fig. 3A, left | linear regression | 0.002<br>0.001 | Female-female, group-housed resident<br>Female-female, single-housed resident |
| Fig. 3A, middle | linear regression | 0.83<br>0.02 | Male-male, group-housed resident<br>Male-male, single-housed resident |
| Fig. 3A, right | linear regression | 0.003<br>0.006 | Male-female, group-housed resident<br>Male-female, single-housed resident |
| Fig. 3B | two-way ANOVA, repeated measures on one factor (A, B, and interaction) | <0.001 <0.001 0.004 | differences in proportion female-female USVs produced during different non-vocal social behaviors |
| | | <0.001 <0.001 <0.001 | differences in proportion male-male USVs produced during different non-vocal social behaviors |
| | | >0.05 <0.001 >0.05 | Differences in proportion male-female USVs |

| | | | |
|---|---|---|---|
| | | | produced during different non-vocal social behaviors |
| Fig. 3B | Post-hoc tests (resident-initiated, visitor-initiated, mutual) | 0.034 0.12 0.05 | differences in proportion female-female USVs produced during different non-vocal social behaviors |
| | | 0.006 0.96<br>0.96 | differences in proportion male-male USVs produced during different non-vocal social behaviors |
